# Desmin intermediate filaments and tubulin detyrosination stabilize growing microtubules in the cardiomyocyte

**DOI:** 10.1101/2021.05.26.445641

**Authors:** Alexander K. Salomon, Sai Aung Phyo, Naima Okami, Julie Heffler, Patrick Robison, Alexey I. Bogush, Benjamin L. Prosser

## Abstract

In heart failure, an increased abundance of post-translationally detyrosinated microtubules stiffens the cardiomyocyte and impedes its contractile function. Detyrosination promotes interactions between microtubules, desmin intermediate filaments and the sarcomere to increase cytoskeletal stiffness, yet the mechanism by which this occurs is unknown. We hypothesized that detyrosination may regulate the growth and shrinkage of dynamic microtubules to facilitate interactions with desmin and the sarcomere. Through a combination of biochemical assays and direct observation of growing microtubule plus-ends in adult cardiomyocytes, we find that desmin is required to stabilize growing microtubules at the sarcomere Z-disk, where desmin also rescue shrinking microtubules from continued depolymerization. Further, reducing detyrosination (tyrosination) promotes frequent depolymerization and inefficient growth of microtubules. This is concomitant with tyrosination promoting the interaction of microtubules with the depolymerizing protein complex of end-binding protein 1 (EB1) and CAP-Gly domain containing linker protein 1 (CLIP1/CLIP170). The futile growth of tyrosinated microtubules reduces their opportunity for stabilizing interactions at the Z-disk, coincident with tyrosination globally reducing microtubule lifetimes and stability. These data provide a model for how intermediate filaments and tubulin detyrosination establish long-lived and physically reinforced microtubules in the cardiomyocyte, and inform on the mechanism of action for therapies that target microtubules for the treatment of cardiac disease.

## Introduction

Microtubules are polymers of α- and β-tubulin that are characterized by cyclical transitions between polymerization and depolymerization, a behavior called dynamic instability[24]. Tuning this dynamic behavior confers unique functionality to specific sub-populations of microtubules[15]. Control of microtubule dynamics is cell type- and context-specific and can occur either by modulating polymer addition or subtraction at the ends, or through lateral interaction with the microtubule[1]. The temporal and spatial control of dynamics can be tuned by post-translational modification of tubulin, which in turn affects the biophysical properties of the microtubule and interactions with microtubule-associated proteins (MAPs)[31]. For example, detyrosination, the post translational removal of the C-terminal tyrosine residue on α-tubulin, has been shown to alter microtubule stability through modulating interactions with multiple effector MAPs [9, 26, 27].

In the cardiomyocyte, microtubules fulfill both canonical roles in intracellular trafficking and organelle positioning[6, 32], as well as non-canonical functions matched to the unique demands of working myocytes. In the interior of the myocyte, microtubules form a predominantly longitudinal network that runs perpendicular to the transverse Z-disks that define the sarcomere, the basic contractile unit of muscle. This microtubule network is required for the delivery of essential cargo in the myocyte, including ion channels and membrane proteins required for muscle excitation, as well as the distribution of RNAs and the translational machinery to maintain and grow new sarcomeres[32, 35]. To perform this role, microtubules must also withstand the high forces and changes in cell geometry inherent to cardiac contraction.

To this end sub-populations of microtubules form physical connections with the Z-disk that serve as lateral reinforcements along the length of the microtubule[30]. Upon stimulation and sarcomere shortening, these physically coupled microtubules buckle at short stereotypical wavelengths between sarcomeres to resist the change in myocyte length[30]. Lateral reinforcement has significant mechanical ramifications, as reinforced microtubules can resist forces 3 orders of magnitude greater than isolated microtubules[4, 34]. This viscoelastic resistance, while modest under normal conditions, becomes particularly problematic in heart failure, where proliferation of coupled microtubules stiffens the cardiomyocyte and impairs myocyte motion[7].

Physical coupling of the microtubule to the sarcomere is tuned by detyrosination. Genetic reduction of detyrosination (i.e. tyrosination) by overexpression of tubulin tyrosine ligase (TTL), the enzyme responsible for ligating the terminal tyrosine residue on detyrosinated tubulin, reduces sarcomeric buckling and the viscoelastic resistance provided by microtubules, increasing the contractility of failing myocytes [7]. Tyrosination status also governs microtubule-dependent mechanotransduction in muscle that regulates downstream second messengers and is implicated in myopathic states [19]. Given its ability to lower stiffness and improve the function of myocytes and myocardium from patients with heart failure[7], tyrosination is under pursuit as a novel therapeutic approach. Yet how detyrosination promotes the interaction of microtubules with the sarcomere remains poorly understood.

Several observations suggest this interaction may be mediated at least in part through desmin intermediate filaments that wrap around the Z-disk. Detyrosination promotes microtubule interaction with intermediate filaments[16, 30], and in the absence of desmin, microtubules are disorganized and detyrosination no longer alters myocyte mechanics[30]. Importantly, a recent publication indicates that intermediate filaments can directly stabilize dynamic microtubules in vitro[33]. However, there has been no investigation into the effect of desmin or detyrosination on the dynamics of cardiac microtubules.

Here, using a combination of genetic manipulations, biochemical assays, and direct live-cell observation of dynamic microtubules, we interrogated the effect of desmin depletion and tubulin tyrosination on microtubule dynamics. We find that desmin spatially organizes microtubule dynamics, conferring local stability to both growing and shrinking microtubules at the sarcomere Z-disk. Additionally, we find that tyrosinated microtubules are more dynamic and prone to shrinkage, a characteristic that precludes their ability to efficiently grow between adjacent sarcomeres and form stabilizing interactions at the Z-disk. These findings provide insight into the fundamental organizing principles of myocyte cytoarchitecture and inform on the mechanism of action for therapeutic strategies that target detyrosinated microtubules.

## Results

### Dynamic microtubules are stabilized at the Z-disk and interact with desmin intermediate filaments

To study the dynamics of growing microtubules in mature cardiomyocytes, we treated adult rat cardiomyocytes with adenovirus containing GFP-labeled End-Binding Protein 3 (GFP-EB3) to directly visualize the plus-end of growing microtubules by time-lapse imaging (**S. Movie 1)**. The dynamic properties of microtubules can be quantified as events that mark their transitions from growing (polymerization) to shrinking (depolymerization) states **(Fig. 1a)**. These events consist of catastrophes (transitions from growth to shrinkage), rescues (transitions from shrinkage to growth), and pauses (neither growth nor shrinkage). Conveniently, GFP-EB3 also provided a fainter, non-specific labeling of the protein-rich Z-disk, enabling us to visualize where dynamic events occurred relative to a sarcomeric marker (**Fig. 1b)**.

**Fig. 1.**
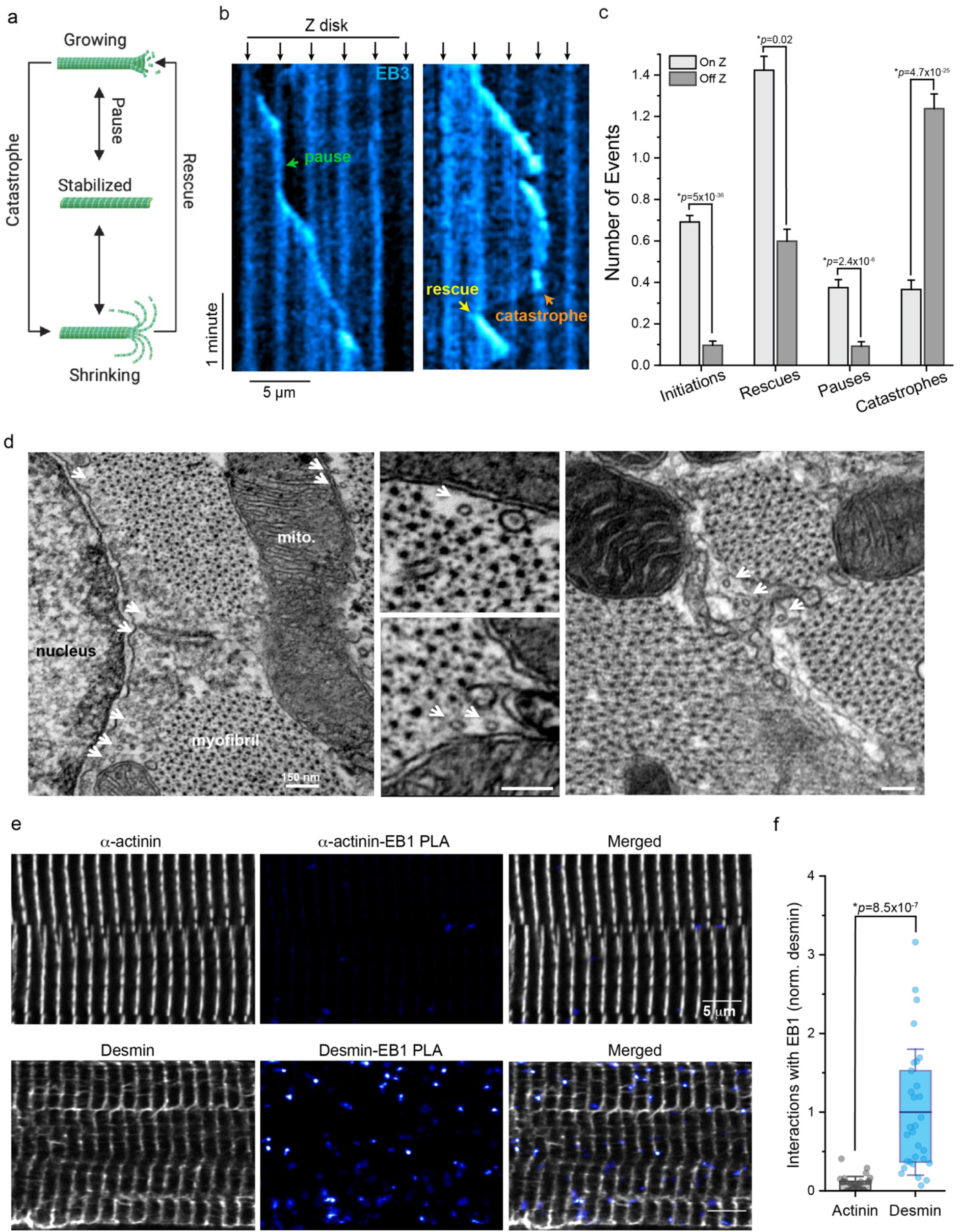
Dynamic microtubules are stabilized at the Z-disk and preferentially interact with desmin intermediate filaments. **(a)** Schematic of the transition states of microtubule dynamics. **(b)** Representative kymograph from control cardiomyocytes transduced with AdV-GFP-EB3; black arrows denote Z-disk and colored arrows denote transition events. **(c)** Quantification of initiation, rescue, pause, and catastrophe events On and Off the Z-disk in control cardiomyocytes (N=19 cells, n=228 events). Bars represent mean ± 1SEM; statistical significance determined with Two Sample Kolmogorov-Smirnov Test. **(d)** Representative EM images from transverse sections of isolated cardiomyocytes. Microtubules are denoted by white arrows. In right hand panel, area between the myofibrils is filled by membranous and filamentous structures consistent with intermediate filaments, which are bisected by microtubules. **(e)** Representative immunofluorescent images & **(f)** quantification of a-actinin-EB1 or Desmin-EB1 PLA interactions in control cardiomyocytes (N=3 rats, n=10 cells per rat). Box represents 25^th^ and 75^th^ percentiles ± 1SD, bolded-line represent mean; statistical significance was determined with Two-sample Student’s T-test.

Under basal conditions, we observed a stark spatial bias in microtubule dynamic behavior, similar to that previously observed[11]. The initiation of microtubule growth, as well as pausing of growth, predominantly occurred on the Z-disk (**Fig. 1c)**. Conversely, catastrophes predominantly occurred off the Z-disk, while rescue from catastrophe again occurred more frequently at the Z-disk. As exemplified in **S. Movies 1-2**, myocyte microtubules tend to grow iteratively from one Z-disk to another, often pausing at each Z-disk region. If a microtubule undergoes catastrophe before reaching a Z-disk, it tends to shrink to a previous Z-disk, where rescue is more likely to occur. These data suggest factors at the Z-disk region strongly bias microtubule behavior and support the initialization and stabilization of growing microtubules.

Electron microscopy images of cardiomyocytes help illustrate the local environment surrounding microtubules at the nanoscale and suggest nearby elements that may stabilize microtubules. As seen in Figure 1d, the microtubules running along the long-axis of the myocyte appear as 25nm diameter tubes coming at the viewer in transverse sections, with a faint halo surrounding them where their C-terminal tails project. Microtubules most commonly run alongside, and not within, the sarcomere-containing myofibrils, squeezing in the gaps between myofibrils and the mitochondria or nucleus. Desmin intermediate filaments also occupy some of these gaps, wrapping around the myofibrils at the level of the Z-disk, and we observe microtubules bisecting through structures that resemble intermediate filaments and which surround the myofibrils at these locations (**Fig. 1d, right)**. To orthogonally probe whether growing microtubules are more likely to interact with the intermediate filament vs. sarcomeric cytoskeleton, we utilized proximity ligation assay (PLA) to probe interactions between the endogenous microtubule plus-end tracking protein end-binding protein 1 (EB1) and either sarcomeric α -actinin or the intermediate filament desmin in adult rat cardiomyocytes. Although α -actinin is the most abundant protein in the Z-disk and expressed at substantially higher levels than desmin[7] (**S. Fig. 1a)**, we observed ∼10-fold more abundant PLA puncta in the desmin-EB1 group compared to α -actinin-EB1, suggesting that the growing end of microtubules are frequently in close proximity to desmin intermediate filaments at the Z-disk.

### Desmin stabilizes growing and shrinking microtubules at the Z-disk

We next directly interrogated the role of desmin in regulating microtubule stability by adenoviral delivery of shRNA to acutely deplete desmin (desmin KD) in cardiomyocytes. Complementing our previous validation of this construct by western blotting[17], we measured a 40-50% reduction in desmin expression after 48 hours of desmin KD (**S. Fig. 1b**). We first interrogated the effect on microtubule stability using a modified subcellular fractionation assay from Fasset et al., 2009[13] that allowed us to separate free tubulin from polymerized tubulin in the dynamic (i.e. cold-sensitive) microtubule pool (**Fig. 2a)**. Acute desmin depletion resulted in an increased free to polymerized ratio in the dynamic microtubule pool (**Fig. 2b**,**c, S. Fig. 1c)**, suggesting that desmin coordinates the stability of dynamic microtubules. We next quantified microtubule acetylation and detyrosination, markers of long-lived microtubules, and found that both were decreased in desmin KD myocytes, without alterations in whole cell tubulin content (**Fig. 2b,c**), suggesting that desmin normally helps maintain microtubule stability.

**Fig. 2.**
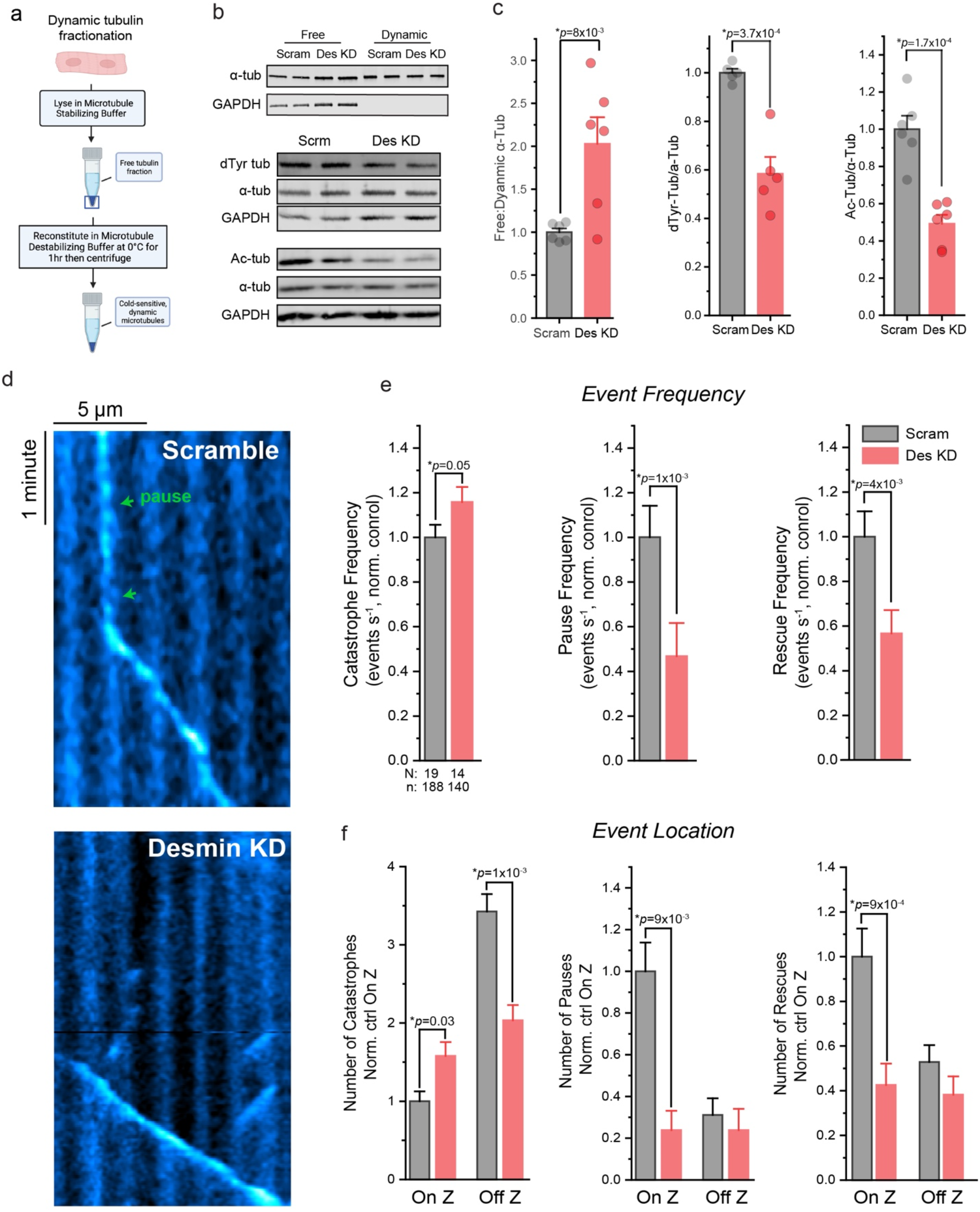
Desmin stabilizes dynamic microtubules at the Z-disk. **(a)** Overview of the cell fractionation assay adapted from Fassett et al.[13] that allows for separation of free tubulin and polymerized microtubules within the dynamic tubulin pool. **(b)** Representative western blot & **(c)** quantification of α-tubulin in free and dynamic microtubule fractions **(top)** or of total dTyr-tubulin, α-tubulin, and acetylated tubulin in whole-cell lysate **(bottom)** from control (Scram) or Desmin knock-down (Des KD) cardiomyocytes (N=3 rats, n=5 WB technical lanes for dtyr and 6 for acetyl and tubulin fractions). **(d)** Representative EB3-GFP kymograph from Scram **(top)** or Des KD **(bottom)** cardiomyocytes. **(e)** Quantification of catastrophe, pause, and rescue event frequencies and **(f)** event locations in Scram or Des KD cardiomyocytes (N=cells, n=events). Bar represents mean ± 1SEM; statistical significance for C was determined with Two-sample Student’s T-test, and for E and F was determined with Two-sample Kolmogorov-Smirnov test.

Next, we directly quantified plus-end microtubule dynamics by EB3-GFP upon desmin depletion. Blind quantification of global event frequency revealed that desmin depletion modestly increased the frequency of catastrophes while more robustly reducing both the frequency of rescues and pauses (**Fig. 2d,e**). As seen in **S. Movies 3-4**, upon desmin depletion (**S. Movies 4)** microtubule growth still initiated at the Z-disk, but the iterative, longitudinal growth from one Z-disk to another seen in control cells (**S. Movies 3)** was lost. Instead, microtubules often grew past Z-disk regions without pausing, and following catastrophe they were less likely to be rescued at the previous Z-disk (**Fig. 2d,f)**. Interrogation of where dynamic events occurred in relation to the Z-disk revealed that desmin depletion specifically increased the number of catastrophes that occurred on the Z-disk, while reducing the number of catastrophes that occurred off the Z-disk (**Fig. 2f**). More strikingly, desmin depletion markedly reduced the number of pauses and rescues that occur specifically on the Z-disk, while not affecting pause or rescue behavior elsewhere (**Fig. 2f**). Together, these results indicate that desmin spatially coordinates microtubule dynamics and stabilizes both the growing and shrinking microtubule at the Z-disk.

Cardiomyocytes from global, desmin germ-line knockout mice are characterized by misaligned and degenerated sarcomeres with a disorganized microtubule network [5, 30]. Gross restructuring of the myofilaments could affect microtubule dynamics due to a change in the physical environment that is permissive to microtubule growth, for example by increasing the spacing between Z-disks of adjacent myofilaments. To assess if our comparatively brief desmin depletion altered myofilament spacing or alignment, we performed quantitative measurements on electron micrographs from desmin KD cardiomyocytes. Blind analysis indicated that this relatively short-term desmin depletion did not detectably alter myofilament spacing or alignment (**S. Fig. 2**), consistent instead with a direct stabilizing effect of desmin intermediate filaments on the microtubule network.

We next interrogated functional consequences of this reduced microtubule stability driven by desmin depletion. As a reduction in detyrosinated microtubules and their association with the Z-disk is associated with reduced cardiomyocyte viscoelasticity[30], we hypothesized that desmin-depleted myocytes would be less stiff. To test this, we performed transverse nanoindentation of cardiomyocytes and quantified Young’s modulus of the myocyte over a range of indentation rates. Desmin depletion specifically reduced the rate-dependent viscoelastic stiffness of the myocyte without significantly altering rate-independent elastic stiffness (**S. Fig. 3a,b)**. Reduced viscoelasticity is consistent with reduced transient interactions between dynamic cytoskeletal filaments.

To directly test if the reduction in desmin alters microtubule buckling between sarcomeres, we performed a semi-automated, blind analysis of microtubule buckling, as in our previous work[30]. In control cells, most microtubules buckle in a clear sinusoidal pattern with a wavelength corresponding to the distance of a contracted sarcomere (∼1.5-1.9 µm) (**S. Fig. 3c,d) (S. Movie 5**). Upon desmin depletion, fewer polymerized microtubules were observed in general, with more chaotic deformations and organization upon contraction (**S. Movie 6**). For microtubules that did buckle, we observed reductions in the amplitude of buckles (**S. Fig. 3d**) and the proportion of microtubules that buckled at wavelengths corresponding to the distance between 1 or 2 sarcomeres (1.5-1.9 or 3.0-3.8 µm, respectively) (**S. Fig. 3e,f)**. Combined, these results are consistent with desmin coordinating the physical tethering and lateral reinforcement of detyrosinated microtubules at the cardiomyocyte Z-disk to regulate myocyte viscoelasticity.

### Tyrosination alters the dynamics of the microtubule network

Next, we sought to determine the effect of detyrosination on the dynamics of the cardiomyocyte microtubule network. To reduce detyrosination, we utilized adenoviral delivery of TTL into isolated adult rat cardiomyocytes[30]. TTL binds and tyrosinates tubulin in a 1:1 complex, and this binding leads to tubulin sequestration. Hence to separate the effects of tubulin tyrosination from tubulin sequestration, we utilized adenoviral delivery of TTL-E331Q (E331Q), a verified catalytically dead mutant of TTL that binds and sequesters tubulin, but does not tyrosinate[8]. We have previously confirmed that TTL overexpression under identical conditions reduces detyrosination below 25% of initial levels, while TTL-E331Q does not significantly affect detyrosination levels with similar overexpression[8]. To specifically quantify the effects of reducing detyrosination on the dynamic microtubule population, we fractionated free and polymerized tubulin as outline above (**Fig. 2a**). Expression of TTL, but not E331Q, resulted in significantly less detyrosinated tubulin in the dynamic microtubule pool (**Fig. 3a**). Further, only TTL expression shifted tubulin away from the polymerized fraction towards the free tubulin fraction, resulting in an increased ratio of free: polymerized tubulin (**Fig. 3a, S. Fig. 4a**). This suggests that tyrosination effects the cycling of tubulin within the dynamic microtubule pool. If indeed tyrosinated microtubules are more dynamic, then levels of acetylation, a canonical marker of long-lived microtubules[36], should also be decreased by TTL. Consistent with this, TTL, but not E331Q, led to a robust reduction in levels of microtubule acetylation, suggesting that tyrosination reduces microtubule lifetime in the cardiomyocyte (**Fig. 3b)**.

**Fig. 3.**
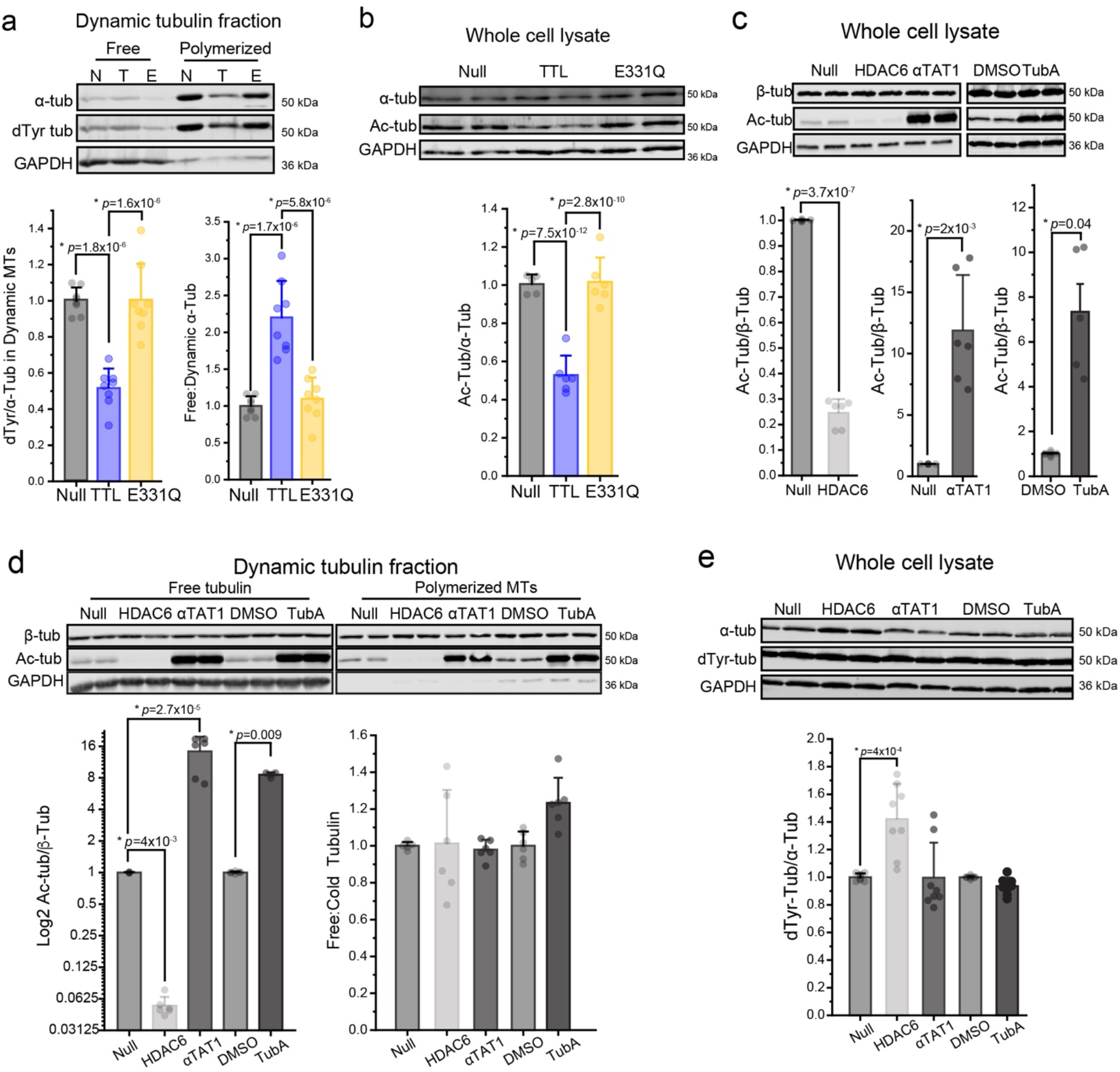
TTL reduces microtubule stability through its tyrosinase activity. **(a)** Representative western blot (**top**) and quantification (**bottom**) of α-tubulin and detyrosinated (dTyr) tubulin in free and cold-sensitive dynamic microtubule fractions from adult rat cardiomyocytes treated with null, TTL, or TTL-E331Q adenoviruses; detyrosinated tubulin values are normalized to α-tubulin in cold-sensitive fraction (N=4 rats, n=8 WB technical lanes). **(b)** Representative western blot **(top)** and quantification **(bottom)** of α-tubulin and acetylated tubulin in whole-cell lysate from null, TTL, or E331Q expressing cardiomyocytes (N=3 rats, n=6 WB technical lanes). **(c)** Validation of HDAC6 and αTAT1 constructs and Tubastatin A (TubA) treatment. Representative western blot **(top)** and quantification **(bottom)** of a-tubulin and acetylated tubulin in whole-cell lysate from adult rat cardiomyocytes treated with null, HDAC6, or αTAT1 adenoviruses, or DMSO or 1 mM TubA treatment overnight (N=3 rats, n=6 WB technical lanes). **(d)** Representative western blot **(top)** and quantification **(bottom)** of α-tubulin and acetylated tubulin, in free and polymerized dynamic fractions. Lysate from cardiomyocytes were infected with null, HDAC6, or αTAT1 adenoviruses, or DMSO or 1 mM TubA overnight (N=3 rats, n=6 WB technical lanes). **(e)** Representative western blot **(top)** and quantification **(bottom)** of α-tubulin and detyrosinated tubulin in whole-cell lysate from adult rat cardiomyocytes treated with null, HDAC6, or αTAT1 adenoviruses, or DMSO or 1 mM TubA treatment overnight (N=4 rats, n=8 WB technical lanes). Bar represents mean ± 1SEM; statistical significance for (a) and (b) was determined with one-way ANOVA with post-hoc test, and for (c) to (e) was determined with Two-sample Student’s T-test.

As acetylation itself is linked to microtubule stability[12, 38], the TTL-dependent change in the dynamic microtubule pool **(Fig. 3b)** could be directly related to tyrosination, or it could be a secondary effect due to the reduction in acetylation. To discriminate between these two hypotheses, we directly modulated acetylation. To this end, we developed adenoviral constructs encoding histone deacetylase 6 (HDAC6) and α tubulin acetyltransferase 1 (αTAT1). HDAC6 expression reduced total microtubule acetylation to 25% of initial levels (**Fig. 3c**) and αTAT1 expression increased acetylation 12-fold (**Fig. 3c**). Because αTAT1 has been shown to modulate microtubule dynamics independent of enzymatic activity[18], we also used a pharmacological inhibitor of HDAC6, Tubastatin A (TubA) to increase acetylation through an orthogonal approach (**Fig. 3c**). Having validated robust tools to modulate acetylation, we next determined the effect of acetylation on the dynamic microtubule pool utilizing the same fractionation assay. Neither increasing or decreasing acetylation altered the free:polymerized tubulin ratio (**Fig. 3d, S. Fig. 4b**). Given that modulating tyrosination altered levels of acetylation (**Fig. 3c**), we also asked whether this relationship was reciprocal. However, whole cell levels of detyrosination were largely unaffected by modulating acetylation (**Fig. 3e**), except for a modest increase with HDAC6 expression that may be related to HDAC6 association with microtubules increasing their stability and availability for detyrosination[2]. Together, these results suggest tyrosination directly alters cardiomyocyte microtubule stability, independent of corresponding changes in acetylation.

### Tyrosination promotes catastrophe of growing microtubules

Next, to precisely quantify the effects of tyrosination on the dynamics of individual microtubules, we overexpressed either Null, TTL, or E331Q viruses in conjunction with GFP-EB3 in adult rat cardiomyocytes. Although EB interaction is thought to be unaffected by microtubule detyrosination[26], we first wanted to validate that EB3 labeling of microtubules did not systematically differ with TTL expression. EB3 fluorescence intensity along the length and at the tip of the microtubule was unchanged in control, TTL or E331Q expressing cells (**S. Fig. 4c**), indicating that EB3 expression or labeling of microtubules was not altered by our experimental interventions.

As seen in **S. Movie 7**, microtubules in TTL expressing cells still initiated growth at the Z-disk, but often had shorter runs and underwent catastrophe prior to reaching a subsequent Z-disk. Consistently, TTL overexpression significantly increased the frequency of catastrophes, while reducing the frequency of pausing (**Fig. 4a,b**). E331Q expression did not alter event frequency compared to control cells **(S. Movie 8)**, suggesting a tyrosination-specific effect on microtubule dynamics (**S. Fig. 4d)**. Further examination of spatial dynamics revealed that the effect of TTL on microtubule breakdown was agnostic to subcellular location; TTL similarly increased the number of catastrophes both on and off the Z-disk. In contrast, TTL reduced the number of pauses specifically on the Z-disk (**Fig. 4c**). As a readout of inefficient growth, TTL increased the tortuosity of microtubule trajectories, defined as the ratio of growth distance to net growth (**Fig. 4d**). Combined, the lack of stabilization at the Z-disk and more frequent catastrophes resulted in tyrosinated microtubules depolymerizing ∼5 fold as often before successfully crossing a Z-disk when compared to either null or E331Q expressing cells (**Fig. 4d**). In sum, this data indicates that tyrosination increases the stochastic transition to microtubule breakdown irrespective of subcellular location, and that tyrosinated microtubules inefficiently navigate successive sarcomeres with fewer stabilizing interactions at the Z-disk.

**Fig. 4.**
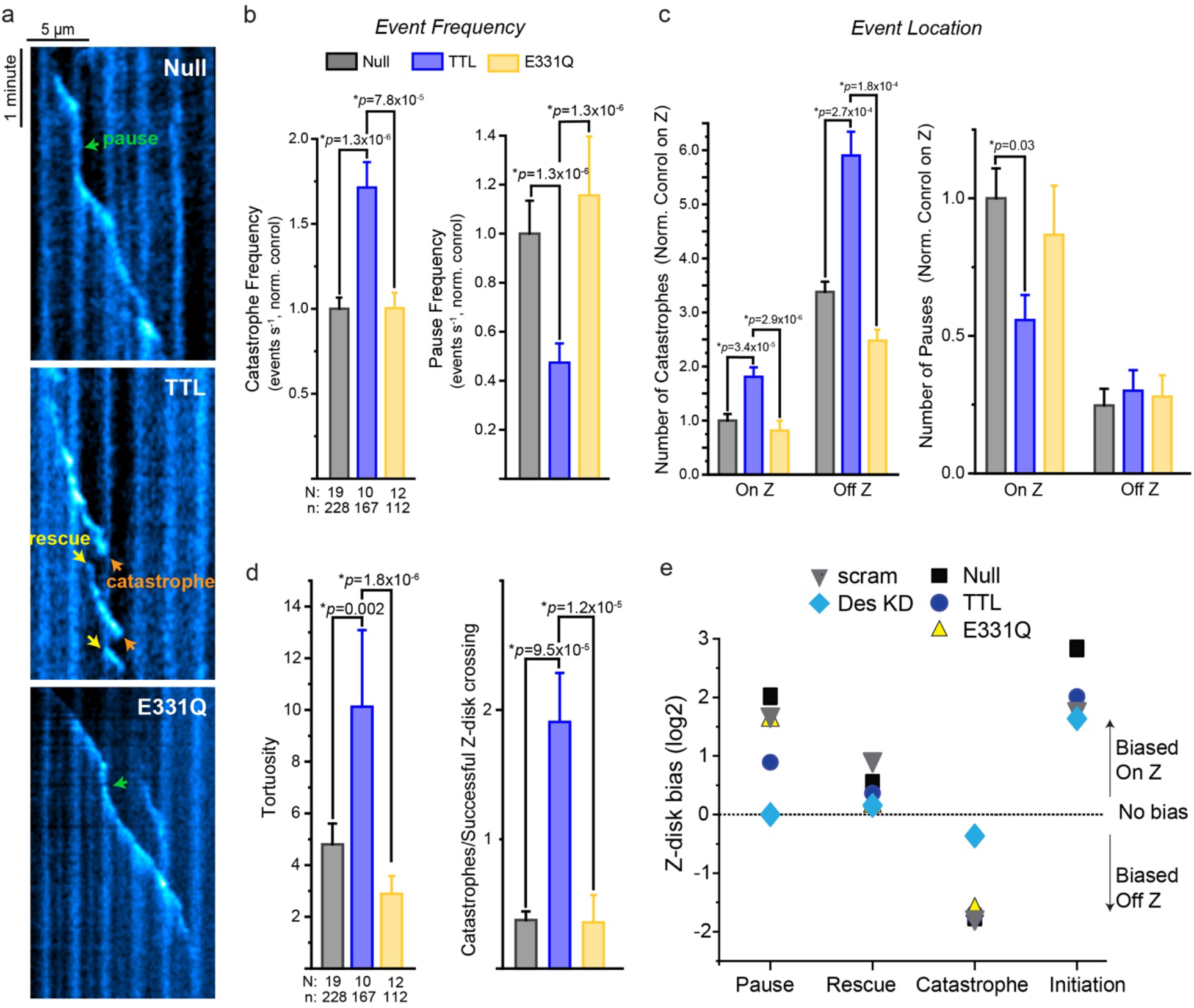
Tyrosinated microtubules are more dynamic. **(a)** Representative kymographs from cardiomyocytes treated with EB3-GFP plus null, TTL or E331Q adenoviruses. **(b)** Quantification of catastrophe and pause event frequencies and **(c)** event locations in cardiomyocytes treated with EB3-GFP plus null, TTL or E331Q adenoviruses (N=cells, n=events). **(d)** Gross measurements of microtubule dynamics. **(left)** Tortuosity, the distance a microtubule grows divided by its displacement, & **(right)** number of catastrophes in relation to number of successful Z-disk crossing in cardiomyocytes treated with EB3-GFP plus null, TTL or E331Q adenoviruses. **(e)** Z-disc bias score (log2 transformation of the ratio of events that occurred On vs. Off the Z-disk) for all experimental conditions. Bar represents mean ± 1SEM; statistical significance was determined with Kruskal-Wallis ANOVA with post-hoc test.

To summarize how our different interventions (tyrosination, desmin depletion) affected the spatial organization of microtubule behavior, we took the ratio of events that occurred on vs. off the Z-disk and performed a log2 transform, calculating a “Z-disk bias” for each type of dynamic event (**Fig. 4e**). Of note, this metric only reflects the spatial bias of events, not their frequencies. TTL reduced the preference for microtubule pausing at the Z-disk, but did not affect the spatial preference of rescues, catastrophes, or initiations. Desmin depletion, on the other hand, virtually eliminated the typical Z-disk bias for pauses, rescues, or fewer catastrophes. Initiations had a strong Z-disk bias regardless of intervention, which likely reflects nucleating events from microtubule organizing centers at Golgi outposts proximal to the Z-disk that are not affected by these manipulations[25].

### Tyrosination increases EB1 and CLIP170 association on microtubules

Next, we wanted to determine why tyrosinated microtubules exhibit increased catastrophe frequencies. Several pieces of evidence suggest that the tyrosinated or detyrosinated status of the microtubule alone is likely insufficient to alter microtubule dynamics[20, 37], but instead the PTM exerts its effect by governing the interaction of stabilizing/destabilizing MAPs with the microtubule [9, 27]. There are two prominent examples of tyrosination altering interactions with depolymerizing effector proteins in the literature. First, mitotic centromere-associated kinesin (MCAK/Kif2C) is a depolymerizing MAP that preferentially binds and depolymerizes tyrosinated microtubules [27]. Second, a recent in vitro reconstitution study indicates that tyrosination promotes the binding of CLIP170 on microtubule plus ends, which synergizes with EB1 to increase the frequency of catastrophe[9]. This mechanism has not been examined in cells. Due to its low abundance in the post-mitotic cardiomyocyte, our attempts to detect and knock down MCAK levels were unreliable; we thus hypothesized that tyrosination may promote the interaction of EB1 and CLIP170 on microtubules to promote their destabilization and catastrophe.

To test this hypothesis, we utilized a PLA to test whether EB1 and CLIP170 interactions on cardiac microtubules were guided by tyrosination. We first performed control assays to ensure the specificity of this PLA assay and ask whether EB1-CLIP170 interactions are observed on intact microtubules. No PLA puncta were observed when primary antibodies against EB1 or CLIP170 were excluded from the PLA assay (**S. Fig. 5a)**. Further, the majority of EB1-CLIP170 interactions co-localized directly on super-resolved microtubules (**Fig. 5a)** indicating that interactions occur primarily on the polymerized microtubule. We next evaluated whether this interaction was sensitive to tyrosination. First, we ensured that global levels of EB1 or CLIP170 were not changing due to TTL or E331Q expression (**Fig. 5b,c)**. We then quantified specific interactions of EB1-CLIP170 that were occurring on microtubules by thresholding the microtubule and PLA images, quantifying the fractional area covered by their overlap, and normalizing that area to the microtubule coverage in the same image plane (**S. Fig. 5b)**. As shown in Figure 5d, TTL increased the number of EB1-CLIP170 interactions per microtubule area by ∼4-fold relative to control or E331Q transduced cardiomyocytes (**Fig. 5d**), despite unchanging levels of EB1 or CLIP170. As this interaction has been demonstrated to be sufficient to robustly increase the catastrophe frequency of dynamic microtubules [9], we conclude that tyrosination destabilizes cardiac microtubules at least in part by promoting increased association with the destabilizing effector complex of EB1 and CLIP170.

**Fig. 5.**
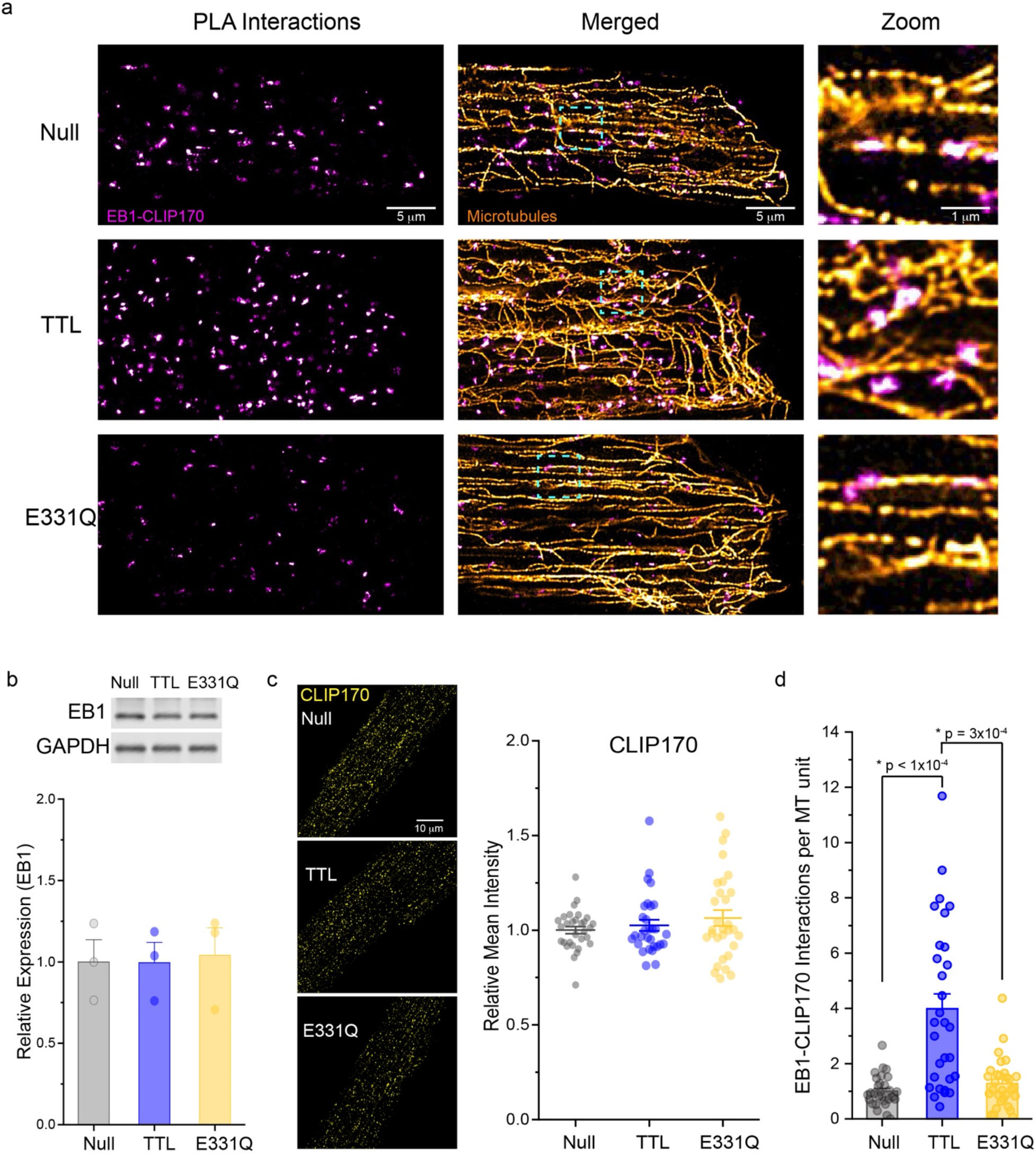
Tyrosination promotes EB1 and CLIP170 interactions on cardiomyocyte microtubules. **(a)** Representative AiryScan Joint Deconvoluted immunofluorescent images of EB1-CLIP170 PLA interactions in adult rat cardiomyocytes treated with null, TTL, or TTL-E331Q adenoviruses. **(b)** Representative western blot **(top)** and quantification **(bottom)** of EB1 in whole-cell lysate from adult rat cardiomyocytes treated with null, TTL, or E331Q adenoviruses for 48h (N=3 rat, n=3 WB technical lanes). **(c)** Representative immunofluorescent images **(left)** and quantification **(right)** of CLIP170 in adult rat cardiomyocytes treated with null, TTL, or E331Q adenoviruses for 48h (N=3 rats, n=10 cells per rat). **(d)** Quantification of EB1-CLIP170 PLA interactions in adult rat cardiomyocytes treated with null, TTL, or TTL-E331Q adenoviruses (N=3 rats, n=10 cells per rat). Bar represents mean ± 1SEM, and middle line in box graph represents mean ± 1SEM; statistical significance for (b) was determined with one-way ANOVA with post-hoc test, and for (c) and (d) was determined with Kruskal-Wallis ANOVA with post-hoc test.

## Discussion

In this paper we identify that 1) desmin intermediate filaments structure and stabilize growing microtubules; 2) microtubule tyrosination promotes destabilizing interactions with EB1+CLIP170; 3) the catastrophe-prone nature of tyrosinated microtubules precludes their ability to faithfully traverse and be stabilized at successive Z-disks. When combined with recent *in vitro* studies using reconstituted microtubules and intermediate filaments[9, 33], our *in cellulo* findings provide a molecular model for how changing levels of desmin and detyrosination may synergistically control cytoskeletal stability in the heart. These findings also inform on the mechanism of action for therapeutic approaches that target the tyrosination cycle for the treatment of heart failure.

This study represents the first direct observation that tyrosination increases the dynamics of cardiac microtubules. A recent report provides compelling evidence to support the long-standing belief that altered dynamicity does not arise from tyrosination/detyrosination itself, but instead through PTM-dependent changes in recruitment of effector proteins [9, 20, 26]. The C-terminal tyrosine on unstructured tubulin tails is likely insufficient to influence lateral contacts between tubulin dimers in the microtubule lattice that confer stability. Yet removal of the large hydrophobic tyrosine residue, and the subsequent exposure of acidic residues, will alter hydrophobic and electrostatic interactions on the outer surface[27] of the polymerized microtubule. Through such a mechanism, tyrosination can promote microtubule dynamics via increased interaction with destabilizing MAPs, or through decreased interaction with stabilizing MAPs.

As case in point, tyrosination increases the affinity of the depolymerizing kinesin MCAK for the microtubule, decreasing microtubule stability[27]. The low abundance of MCAK in the cardiomyocyte, while not ruling out a physiological rule, motivated interrogation into alternative stabilizing or destabilizing effector proteins. Tyrosination is also known to impact the recruitment of plus-end tip proteins (+Tips), such as CLIP170 and p150 glued[26], which can tune microtubule dynamics through either direct or indirect effects. +Tip proteins can couple the growing microtubule plus end to subcellular targets through a search and capture mechanism, where dynamic, probing microtubules ‘search’ for interacting sites on the plasma membrane, chromosomes, and organelles which are ‘captured’ via +Tip proteins[21, 23]. While +Tip interaction with a target often stabilizes searching microtubules, Chen et al. recently found that the tyrosination-dependent recruitment of the +Tip protein CLIP170 to growing microtubules paradoxically led to a synergistic interaction with EB1 that selectively reduced the stability of tyrosinated microtubules, increasing their catastrophe frequency. Here we find that in cardiomyocytes, while tyrosination has no effect on the global levels of either EB1 or CLIP170 (**Fig. 5b,c**), it robustly increases the frequency of their interaction on microtubules (**Fig. 5c,d)**, concomitant with increased frequency of catastrophe (**Fig. 4b**). While this does not rule out other potentially destabilizing effects of tyrosination, it provides one mechanism for the increased dynamicity/decreased stability of tyrosinated microtubules.

We also identified that the intermediate filament desmin provides structure to the growing microtubule network by stabilizing both growing and shrinking microtubules at the cardiomyocyte Z-disk. What is the mechanism of desmin-dependent stabilization? A recent elegant *in vitro* study using reconstituted vimentin intermediate filaments and microtubules indicates that intermediate filaments are sufficient to stabilize growing microtubules through electrostatic and hydrophobic interactions[33]. Dynamic microtubules interacting with intermediate filaments reduces catastrophes and promotes rescues, in strong accordance with our *in cellulo* findings here. While MAPs may also be involved in modulating microtubule-intermediate filament interactions, this direct effect is sufficient to explain the primary phenotypes observed upon desmin depletion (i.e. increased catastrophes and reduced pausing in the absence of desmin at the Z-disk, and a loss of rescues at the Z-disk).

The intermediate filament stabilization of growing microtubules would then provide a longer-lived microtubule substrate to facilitate reinforcing, detyrosination-dependent interactions, such as those previously documented between desmin and the microtubule through intermediates such as Kinesin-1[22] or members of the plakin family of scaffolding proteins[14]. Desmin-mediated frictional interaction along the length of the microtubule may also lead to the loss of tubulin dimers at sites of contact; these lattice defects are replaced by GTP-tubulin, which upon microtubule catastrophe may function as a rescue site[3]. Lateral interactions between microtubules and intermediate filaments govern microtubule mechanical behavior upon compressive loading of microtubules[34] allowing desmin to orchestrate microtubule buckling in the cardiomyocyte.

Combined with past and current work, we propose a unifying model for microtubule-intermediate filament interactions in the cardiomyocyte and how they contribute to myocardial mechanics (**Fig. 6**). Detyrosinated microtubules, with less frequent depolymerization, experience more chance interactions with intermediate filaments at the Z-disk. The altered surface chemistry of detyrosinated microtubules may also strengthen the electrostatic interactions with intermediate filaments and additional cross-linking proteins. The periodic, lateral reinforcement of microtubules increases their stability, leading to longer-lived microtubules and providing a dynamic cross-link with the sarcomere, increasing the viscoelastic resistance to myocyte motion and the ability of microtubules to bear and transduce mechanical stress. Increased microtubule lifetimes also promote microtubule acetylation, which itself increases the ability of microtubules to withstand mechanical stress[28] and increases myocyte viscoelasticity[10]. In the setting of heart disease, the increased abundance of both desmin intermediate filaments and detyrosinated microtubules thus promotes a feed-forward substrate for enhanced mechanotransduction and myocardial stiffening. Therapeutic strategies that selectively re-tyrosinate the network – independent of grossly depolymerizing microtubules – may thus reduce myocardial stiffening via restoring dynamicity to cardiac microtubules.

**Fig. 6.**
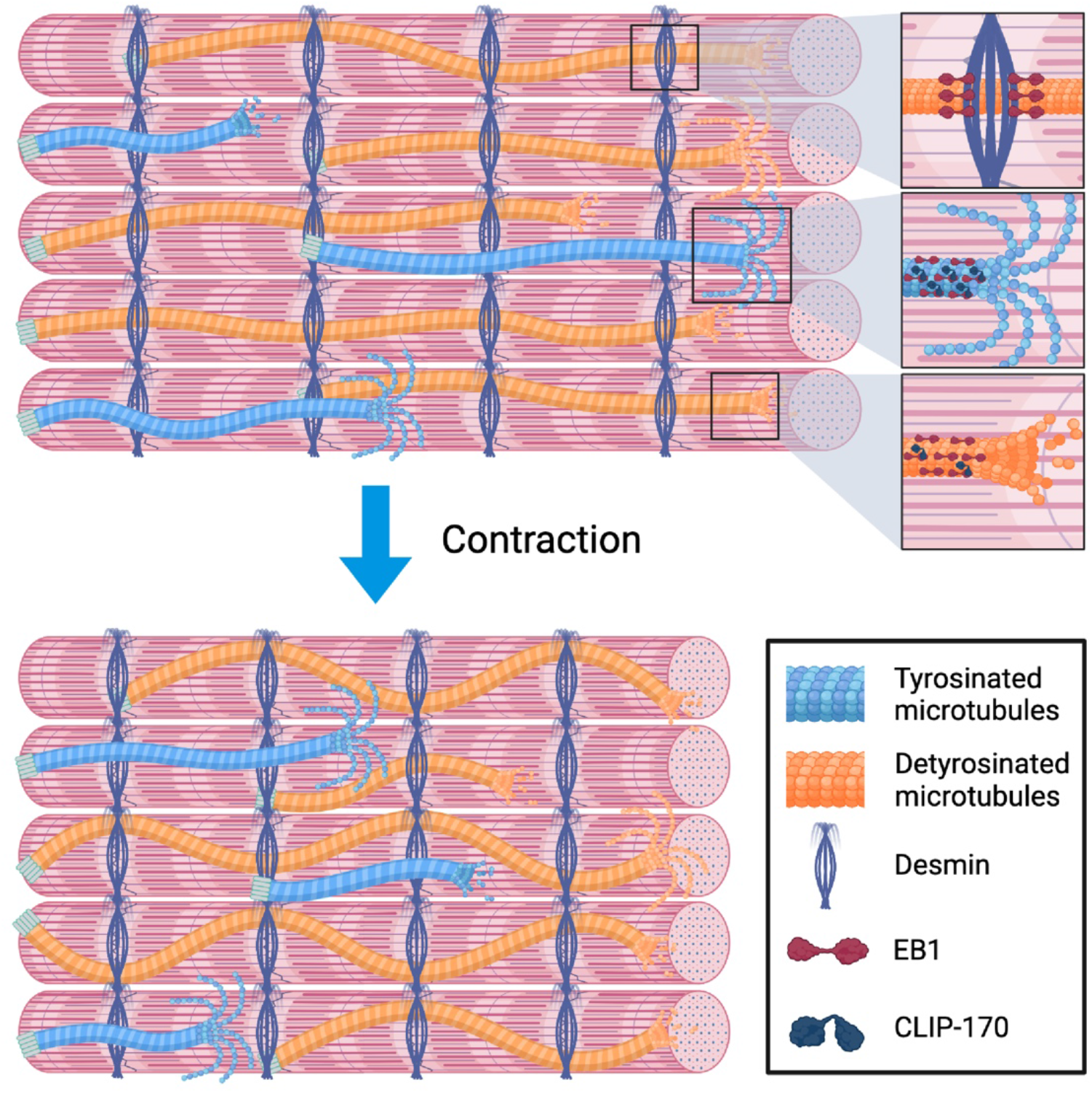
Cartoon summary of the results: Desmin intermediate filaments and tubulin detyrosination stabilize growing microtubules in the cardiomyocyte.

## Methods

### Animals

Animal care and procedures were approved and performed in accordance with the standards set forth by the University of Pennsylvania Institutional Animal Care and Use Committee and the Guide for the Care and Use of Laboratory Animals published by the US National institutes of Health.

### Rat cardiomyocyte isolation and culture

Primary adult ventricular myocytes were isolated from 6-to 8-week-old Sprague Dawley rats as previously described[29]. Briefly, rats were anesthetized under isoflurane while the heart was removed and retrograde perfused on a Lutgendorf apparatus with a collagenase solution. The digested heart was then minced and triturated using a glass pipette. The resulting supernatant was separated and centrifuged at 300 revolutions per minute to isolate cardiomyocytes that were resuspended in rat cardiomyocyte media at a density that ensured adjacent cardiomyocytes did not touch. Cardiomyocytes were cultured at 37°C and 5% CO_2_ with 25 μmol/L of cytochalasin D. The viability of rat cardiomyocytes upon isolation was typically on the order of 50-75% rod-shaped, electrically excitable cells, and the survivability for 48hrs of culture is >80% (See Heffler et al. [17]for our quantification of cardiomyocyte morphology in culture).

Rat cardiomyocyte media: medium 199 (Thermo Fisher 115090) supplemented with 1x Insulin-transferrin-selenium-X (Gibco 51500056), 1 μg μl^-1^ primocin (Invivogen ant-pm-1), 20 mmol/L HEPES at pH 7.4 and 25 μmol/L cytochalasin D.

### Fractionation assay of free tubulin and cold-sensitive microtubules

Free tubulin was separated from cold-labile microtubules using a protocol adapted from Tsutusi et al., 1993 and Ostlud et al., 1979. Isolated rat cardiomyocytes were washed once with PBS and homogenized with 250 ul of microtubule stabilizing buffer using a tissue homogenizer. The homogenate was centrifuged at 100,000 xg for 15 minutes at 25°C and the resulting supernatant was stored at -80°C as the free tubulin fraction. The pellet was resuspended in ice-cold microtubule destabilizing buffer and incubated at 0°C for 1 hour. After centrifugation at 100,000 xg for 15 minutes at 4°C the supernatant containing the cold-labile microtubule fraction was stored at -80°C.

Microtubule stabilizing buffer: 0.5 mM MgCl_2_, 0.5 mM EGTA, 10mM Na_3_PO_4_, 0.5 mM GTP, and 1X protease and phosphatase inhibitor cocktail (Cell Signaling #5872S) at pH 6.95

Microtubule destabilizing buffer: 0.25 M sucrose, 0.5 mM MgCl_2_ 10 mM Na_3_PO_4_, 0.5 mM GTP, and 1X protease and phosphatase inhibitor cocktail (Cell Signaling #5872S) at pH 6.95

### Western blot

For whole cell protein extraction, isolated rat cardiomyocytes were lysed in RIPA buffer (Cayman #10010263) supplemented with protease and phosphatase Inhibitor cocktail (Cell Signaling #5872S) on ice for 1 hour. The supernatant was collected and combined with 4X loading dye (Li-COR #928-40004), supplemented with 10% 2-mercaptoethonol, and boiled for 10 minutes. The resulting lysate was resolved on SDS-PAGE gel and protein was blotted to nitrocellulose membrane (Li-COR #926-31902) with mini Trans-Blot Cell (Bio-Rad). Membranes were blocked for an hour in Odyssey Blocking Buffer (TBS) (LI-COR #927-50000) and probed with corresponding primary antibodies overnight at 4 °C. Membranes were rinsed with TBS containing 0.5% Tween 20 (TBST) three times and incubated with secondary antibodies TBS supplemented with extra 0.2% Tween 20 for 1 hour at room temperature. Membranes were washed again with TBST (0.5% Tween 20) and imaged on an Odyssey Imager. Image analysis was performed using Image Studio Lite software (LI-COR). All samples were run in duplicates and analyzed in reference to GAPDH.

### Antibodies and labels

Acetylated tubulin; mouse monoclonal (Sigma T6793-100UL); western blot: 1: 1000

Detyrosinated tubulin; rabbit polyclonal (Abcam ab48389); western blot: 1: 1000

Alpha tubulin; mouse monoclonal, clone DM1A (Cell Signaling #3873); western blot: 1:1000

Alpha tubulin; mouse monoclonal, clone DM1A conjugated to AlexaFluor (AF) 488 (Cell Signaling #8058S); immunofluorescence: 1:100

Beta tubulin; rabbit polyclonal (Abcam ab6046); western blot: 1:1000

Tyrosinated tubulin; mouse monoclonal (Sigma T9028-.2ML); immunofluorescence: 1:1000

Anti-sarcomeric alpha actinin; mouse monoclonal, clone EA-53 (Abcam ab9465); western blot, PLA: 1:1000

Desmin; rabbit polyclonal (ThermoFisher PA5-16705); western blot, immunofluorescence: 1: 1000

Desmin; mouse monoclonal, clone D33 (Agilent Technologies M076029-2); western blot, PLA; 1:500

EB1; rabbit polyclonal (Sigma E3406-200UL); western blot, PLA: 1:400

CLIP170; mouse monoclonal, clone F-3 (Santa Cruz sc-28325); immunofluorescence, PLA: 1:100

GAPDH; mouse monoclonal (VWR GenScript A01622-40); western blot: 1:1000

Goat anti-mouse AF 488 (Life Technologies A11001); immunofluorescence: 1:1000

Goat anti-rabbit AF 565 (Life Technologies A11011); immunofluorescence: 1:1000

IRDye 680RD Donkey anti-Mouse IgG (H + L) (LI-COR 926-68072); western blot: 1:10000

IRDye 800CW Donkey anti-Rabbit IgG (H + L) (LI-COR 926-32213); western blot: 1:10000

Duolink In Situ PLA probe Anti-Rabbit PLUS, Donkey anti-Rabbit IgG (H + L) (Sigma DUO92002); PLA: 1:5 (as per manufacturer’s protocol)

Duolink In Situ PLA probe Anti-Mouse MINUS, Donkey anti-Mouse IgG (H + L) (Sigma DUO92004); PLA: 1:5 (as per manufacturer’s protocol)

### Microtubule Dynamics by EB3

Isolated rat cardiomyocytes were infected with an adenovirus containing an EB3-GFP construct. After 48 hours, cells were imaged on an LSM Zeiss 880 inverted Airyscan confocal microscope using a 40X oil 1.4 numerical aperture objective. Cells expressing EB3-GFP only at the tip were imaged for four minutes at a rate of 1fps. Files were blinded, Gaussian blurred, and Z-compressed using Image J (National Institutes of Health) to generate kymographs. The number of catastrophes, rescues, and pauses were recorded per kymograph in addition to manual tracing of microtubule runs to quantify time, distance, and velocity of microtubule growth or shrinkage. We refer to the entire kymograph as the microtubule ‘track’ that is made up of individual growth and shrinkage events we call ‘runs’. Catastrophe and rescue frequency were calculated per cell by dividing the number of catastrophes or rescues by total time spent in growth or shrinkage time, respectively. Catastrophes and rescues occurring specifically on or off the Z-disk were normalized by the total time of microtubule growth and shrinkage. Experimental values were normalized to their respective control cells (Null for TTL and E331Q, or shScrm for shDes) acquired from the same animals. A minimum of 3 separate cell isolations were performed for each group.

### Immunofluorescence

To stain for desmin: cardiomyocytes were fixed in pre-chilled 100% methanol for 8 minutes at -20°C. Cells were washed 4x then blocked with Sea Block Blocking Buffer (abcam #166951) for at least 1 hour followed by antibody incubation in Sea Block for 24-48 hours. Incubation was followed by washing 3x with Sea Block, then incubated with secondary antibody for 1 hour at RT. Fixed cells were mounted using Prolong Diamond (Thermo #P36961).

To stain for CLIP170: cardiomyocytes were glued to cleaned coverglass (Electron Microscropy Sciences 72222-01) using MyoTak (IonOptix). The cardiomyocytes on coverslips were fixed in 4% paraformaldehyde (Electron Microscropy Sciences 15710) for 10min at RT, followed by 2 washes in PBS, and then permeabilized using 0.25% Triton in PBS for 10min at RT. Cells were washed 3x then blocked with Sea Block Blocking Buffer for at least 1 hour followed by antibody incubation in Sea Block for 24. Incubation was followed by washing 3x with PBS, then incubated with secondary antibody in Sea Block for 1 hour at RT. Fixed cells were mounted using Prolong Diamond.

We used ImageJ to calculate the percent area fraction of desmin or the mean integrated density of CLIP170. An ROI was drawn to include the entire cell boundary. To calculate % area for desmin, we identified the percent fractional coverage of a fluorescence signal over a manually identified threshold for each image as described previously[7]. The mean integrated density data for CLIP170 was collected directly from ImageJ output using unthresholded max-intensity projected images (3 images per cell) of individual cells.

### Buckling analysis

Adult rat cardiomyocytes were isolated as previously described and infected with adenovirus carrying the microtubule-binding protein EMTB chimerically fused to 3 copies of GFP. The purpose of this construct was to label microtubules fluorescently for imaging. The cells were allowed 48 hours to express the construct. All cells chosen were those that contained sufficient brightness and contrast to observe microtubule elements and where the health of the myocyte was not compromised. To interrogate microtubule buckling amplitude and wavelength, cells were induced to contract at 1 Hz 25 V and imaged during the contraction. For analysis, images were blinded, and a microtubule was located that could be followed during the contraction. The backbone was manually traced at rest and during its peak of contraction and the ROI was saved. The ROI was then analyzed using a macro that rotated so that the ROI had the peak of contraction 90 degrees to the axis of contraction to protect from aliasing errors. The program then calculated the distance between the axis of the ROI and its peak and calculated the peak (amplitude) and the width (half wavelength).

### Electron Microscopy

Transmission electron microscopy images were collected as previously described [17]. Images at 7500x were rotated so the cells were parallel to the longitudinal axis. ROIs were generated between adjacent Z-disks to quantify sarcomere spacing and the angle relative to 90°.

### Proximity Ligation Assay (PLA)

Freshly isolated rat cardiomyocytes were untreated or treated for 48 hours with Null, TTL, or E331Q adenoviruses at 37°C with 5% CO_2_. Once viral construct expressions were confirmed using the tagged mCherry, the cardiomyocytes were glued to cleaned coverglass (EMS 72222-01) using MyoTak (IonOptix). The cardiomyocytes on coverslips were fixed in 4% paraformaldehyde for 10min at RT, followed by 2 washes in PBS, and then permeabilized using 0.25% Triton in PBS for 10min at RT. The samples were blocked in Sea Block for 1hour at RT and stored in 4°C until further processing.

The samples were incubated with EB1 and CLIP170 or alpha-actinin or desmin, primary antibodies overnight at 4°C; the coverslips were then washed in PBS for 15min at RT. Following immediately, PLA was performed in humidified chambers using ThermoFisher DuoLink manufacturer’s protocol starting with “Duolink PLA Probe Incubation.” Briefly, the samples were incubated with Duolink PLA secondary antibodies followed by ligation and amplification. Amplification was performed using Duolink FarRed detection reagents (Sigma DUO92013). Post-amplified samples were washed and incubated with alpha-tubulin antibody (DM1A) conjugated to AF 488 (Cell Signaling #8058S) in Sea Block overnight at RT. The processed samples were washed twice with PBS, and the coverslips were mounted using ProLong Diamond Antifade Mountant (Thermo Fisher P36961).

Imaging was performed using Zeiss AiryScan microscope. 6 imaging slices of 0.18mm thickness was sampled for each cell; 10 cells were sampled per group per experiment (N=3, n=30). ImageJ was used to analyze the images. Microtubules and PLA channels were thresholded and the thresholded images were used to construct microtubule-PLA overlap image. An ROI was drawn to outline the cardiomyocyte border. The raw integrated intensities of the thresholded microtubule only, PLA only, and the microtubule-PLA overlap images for each imaging slice was collected. The microtubule-PLA overlap was then normalized to microtubule only to account for cellular and sub-cellular heterogeneity of microtubule density. The average microtubule-normalized microtubule-PLA overlap for one cell was calculated and the data set was constructed by normalizing all values from one experiment to the average control value of that experiment.

### Statistics

Statistical analysis was performed using OriginPro (Version 2018 & 2019). Normality was determined by Shapiro-Wilk test. For normally distributed data, Two-sample Student’s T-test or one-way ANOVA with post-hoc test was utilized as appropriate. For non-normally distributed data, Two-sample Kolmogrov-Smirnov test or Kruskal-Wallis ANOVA was utilized as appropriate. Specific statistical tests and information of biological and technical replicates can be found in the figure legends. Unless otherwise noted, ‘N’ indicates the number of cells analyzed and ‘n’ indicates number of microtubule runs.

## Supporting information

All supplemental figures

EB3 dynamics control 1

EB3 dynamics control 2

EB3 dynamics scramble

EB3 dynamics desmin knockdown

MT buckling scramble

MT buckling desmin knockdown

EB3 dynamics TTL

EB3 dynamics E331Q

## Acknowledgements

The authors thank Matthew Caporizzo for providing the script used to quantify IF fractional coverage, Tim McKinsey for the HDAC6 and αTAT1 constructs, and Keita Uchida for assisting with PLA image analysis. Funding for this work was provided by the National Institute of Health (NIH) R01s-HL133080 and HL149891 to B. Prosser, by the Fondation Leducq Research Grant no. 20CVD01 to B. Prosser, and by the Center for Engineering Mechanobiology to B. Prosser through a grant from the National Science Foundation’s Science and Technology program: 15-48571.

## Author Contributions

Conceptualization: Alexander K. Salomon, Sai Aung Phyo, Naima Okami, Benjamin L. Prosser; Methodology: Alexander K. Salomon, Sai Aung Phyo, Patrick Robision; Formal analysis and investigation: Alexander K. Salomon, Sai Aung Phyo, Naima Okami, Julie Heffler, Patrick Robison, Alexey I. Bogush; Writing – original draft preparation: Alexander K. Salomon, Sai Aung Phyo, Naima Okami, Benjamin L. Prosser; Writing – review and editing: Alexander K. Salomon, Sai Aung Phyo, Naima Okami, Julie Heffler, Patrick Robison, Alexey I. Bogush, Benjamin L. Prosser; Funding acquisition: Benjamin L. Prosser; Supervision: Benjamin L. Prosser

## Abbreviations

MAP: Microtubule-associated protein
PTM: Post-translational modification
MCAK: Mitotic centromere-associated kinesin
TTL: Tubulin tyrosine ligase
E331Q: Catalytically dead TTL
αTAT1: Alpha-tubulin acetyltransferase 1
HDAC6: Histone deacetylase 6
PLA: Proximity ligation assay
EB1: End-binding protein 1
EB3: End-binding protein 3
CLIP170: CAP-Gly domain containing linker protein 1

## References

1. Akhmanova A, Steinmetz MO (2015) Control of microtubule organization and dynamics: two ends in the limelight. Nat Rev Mol Cell Bio 16:711–726. doi: 10.1038/nrm4084

2. Asthana J, Kapoor S, Mohan R, Panda D (2013) Inhibition of HDAC6 Deacetylase Activity Increases Its Binding with Microtubules and Suppresses Microtubule Dynamic Instability in MCF-7 Cells*. J Biol Chem 288:22516– 22526. doi: 10.1074/jbc.m113.489328

3. Aumeier C, Schaedel L, Gaillard J, John K, Blanchoin L, Théry M (2016) Self-repair promotes microtubule rescue. Nat Cell Biol 18:1054–1064. doi: 10.1038/ncb3406

4. Brangwynne CP, MacKintosh FC, Kumar S, Geisse NA, Talbot J, Mahadevan L, Parker KK, Ingber DE, Weitz DA (2006) Microtubules can bear enhanced compressive loads in living cells because of lateral reinforcement. J Cell Biology 173:733–741. doi: 10.1083/jcb.200601060

5. Brodehl A, Gaertner-Rommel A, Milting H (2018) Molecular insights into cardiomyopathies associated with desmin (DES) mutations. Biophysical Rev 10:983–1006. doi: 10.1007/s12551-018-0429-0

6. Caporizzo MA, Prosser BL (2022) The microtubule cytoskeleton in cardiac mechanics and heart failure. Nat Rev Cardiol 19:364–378. doi: 10.1038/s41569-022-00692-y

7. Chen CY, Caporizzo MA, Bedi K, Vite A, Bogush AI, Robison P, Heffler JG, Salomon AK, Kelly NA, Babu A, Morley MP, Margulies KB, Prosser BL (2018) Suppression of detyrosinated microtubules improves cardiomyocyte function in human heart failure. Nat Med 24:1225–1233. doi: 10.1038/s41591-018-0046-2

8. Chen CY, Salomon AK, Caporizzo MA, Curry S, Kelly NA, Bedi KC, Bogush AI, Krämer E, Schlossarek S, Janiak P, Moutin M-J, Carrier L, Margulies KB, Prosser BL (2020) Depletion of Vasohibin 1 Speeds Contraction and Relaxation in Failing Human Cardiomyocytes. Circ Res 127:e14–e27. doi: 10.1161/circresaha.119.315947

9. Chen J, Kholina E, Szyk A, Fedorov VA, Kovalenko I, Gudimchuk N, Roll-Mecak A (2021) α-tubulin tail modifications regulate microtubule stability through selective effector recruitment, not changes in intrinsic polymer dynamics. Dev Cell. doi: 10.1016/j.devcel.2021.05.005

10. Coleman AK, Joca HC, Shi G, Lederer WJ, Ward CW (2021) Tubulin acetylation increases cytoskeletal stiffness to regulate mechanotransduction in striated muscle. J Gen Physiol 153:e202012743. doi: 10.1085/jgp.202012743

11. Drum BML, Yuan C, Li L, Liu Q, Wordeman L, Santana LF (2016) Oxidative stress decreases microtubule growth and stability in ventricular myocytes. J Mol Cell Cardiol 93:32–43. doi: 10.1016/j.yjmcc.2016.02.012

12. Eshun-Wilson L, Zhang R, Portran D, Nachury MV, Toso DB, Löhr T, Vendruscolo M, Bonomi M, Fraser JS, Nogales E (2019) Effects of α-tubulin acetylation on microtubule structure and stability. Proc National Acad Sci 116:201900441. doi: 10.1073/pnas.1900441116

13. Fassett JT, Xu X, Hu X, Zhu G, French J, Chen Y, Bache RJ (2009) Adenosine regulation of microtubule dynamics in cardiac hypertrophy. Am J Physiol-heart C 297:H523–H532. doi: 10.1152/ajpheart.00462.2009

14. Favre B, Schneider Y, Lingasamy P, Bouameur J-E, Begré N, Gontier Y, Steiner-Champliaud M-F, Frias MA, Borradori L, Fontao L (2011) Plectin interacts with the rod domain of type III intermediate filament proteins desmin and vimentin. Eur J Cell Biol 90:390–400. doi: 10.1016/j.ejcb.2010.11.013

15. Forges H de, Bouissou A, Perez F (2012) Interplay between microtubule dynamics and intracellular organization. Int J Biochem Cell Biology 44:266–274. doi: 10.1016/j.biocel.2011.11.009

16. Gurland G, Gundersen GG (1995) Stable, detyrosinated microtubules function to localize vimentin intermediate filaments in fibroblasts. J Cell Biology 131:1275–1290. doi: 10.1083/jcb.131.5.1275

17. Heffler J, Shah PP, Robison P, Phyo S, Veliz K, Uchida K, Bogush A, Rhoades J, Jain R, Prosser BL (2020) A Balance Between Intermediate Filaments and Microtubules Maintains Nuclear Architecture in the Cardiomyocyte. Circ Res 126:e10–e26. doi: 10.1161/circresaha.119.315582

18. Kalebic N, Martinez C, Perlas E, Hublitz P, Bilbao-Cortes D, Fiedorczuk K, Andolfo A, Heppenstall PA (2013) Tubulin Acetyltransferase αTAT1 Destabilizes Microtubules Independently of Its Acetylation Activity. Mol Cell Biol 33:1114–1123. doi: 10.1128/mcb.01044-12

19. Kerr JP, Robison P, Shi G, Bogush AI, Kempema AM, Hexum JK, Becerra N, Harki DA, Martin SS, Raiteri R, Prosser BL, Ward CW (2015) Detyrosinated microtubules modulate mechanotransduction in heart and skeletal muscle. Nat Commun 6:8526. doi: 10.1038/ncomms9526

20. Khawaja S, Gundersen GG, Bulinski JC (1988) Enhanced stability of microtubules enriched in detyrosinated tubulin is not a direct function of detyrosination level. J Cell Biology 106:141–149. doi: 10.1083/jcb.106.1.141

21. Kumar P, Wittmann T (2012) +TIPs: SxIPping along microtubule ends. Trends Cell Biol 22:418–428. doi: 10.1016/j.tcb.2012.05.005

22. Liao G, Gundersen GG (1998) Kinesin Is a Candidate for Cross-bridging Microtubules and Intermediate Filaments SELECTIVE BINDING OF KINESIN TO DETYROSINATED TUBULIN AND VIMENTIN*. J Biol Chem 273:9797–9803. doi: 10.1074/jbc.273.16.9797

23. Mimori-Kiyosue Y, Tsukita S (2003) “Search-and-Capture” of Microtubules through Plus-End-Binding Proteins (+TIPs). J Biochem 134:321–326. doi: 10.1093/jb/mvg148

24. Mitchison T, Kirschner M (1984) Dynamic instability of microtubule growth. Nature 312:237–242. doi: 10.1038/312237a0

25. Oddoux S, Zaal KJ, Tate V, Kenea A, Nandkeolyar SA, Reid E, Liu W, Ralston E (2013) Microtubules that form the stationary lattice of muscle fibers are dynamic and nucleated at Golgi elementsSkeletal muscle microtubule dynamics. J Cell Biology 203:205–213. doi: 10.1083/jcb.201304063

26. Peris L, Thery M, Fauré J, Saoudi Y, Lafanechère L, Chilton JK, Gordon-Weeks P, Galjart N, Bornens M, Wordeman L, Wehland J, Andrieux A, Job D (2006) Tubulin tyrosination is a major factor affecting the recruitment of CAP-Gly proteins at microtubule plus ends. J Cell Biology 174:839–849. doi: 10.1083/jcb.200512058

27. Peris L, Wagenbach M, Lafanechère L, Brocard J, Moore AT, Kozielski F, Job D, Wordeman L, Andrieux A (2009) Motor-dependent microtubule disassembly driven by tubulin tyrosination. J Cell Biol 185:1159–1166. doi: 10.1083/jcb.200902142

28. Portran D, Schaedel L, Xu Z, Théry M, Nachury MV (2017) Tubulin acetylation protects long-lived microtubules against mechanical ageing. Nat Cell Biol 19:391–398. doi: 10.1038/ncb3481

29. Prosser BL, Ward CW, Lederer WJ (2011) X-ROS Signaling: Rapid Mechano-Chemo Transduction in Heart. Science 333:1440–1445. doi: 10.1126/science.1202768

30. Robison P, Caporizzo MA, Ahmadzadeh H, Bogush AI, Chen CY, Margulies KB, Shenoy VB, Prosser BL (2016) Detyrosinated microtubules buckle and bear load in contracting cardiomyocytes. Science 352:aaf0659. doi: 10.1126/science.aaf0659

31. Roll-Mecak A (2019) How cells exploit tubulin diversity to build functional cellular microtubule mosaics. Curr Opin Cell Biol 56:102–108. doi: 10.1016/j.ceb.2018.10.009

32. Scarborough EA, Uchida K, Vogel M, Erlitzki N, Iyer M, Phyo SA, Bogush A, Kehat I, Prosser BL (2021) Microtubules orchestrate local translation to enable cardiac growth. Nat Commun 12:1547. doi: 10.1038/s41467-021-21685-4

33. Schaedel L, Lorenz C, Schepers AV, Klumpp S, Köster S (2021) Vimentin intermediate filaments stabilize dynamic microtubules by direct interactions. Nat Commun 12:3799. doi: 10.1038/s41467-021-23523-z

34. Soheilypour M, Peyro M, Peter SJ, Mofrad MRK (2015) Buckling Behavior of Individual and Bundled Microtubules. Biophys J 108:1718–1726. doi: 10.1016/j.bpj.2015.01.030

35. Uchida K, Scarborough EA, Prosser BL (2021) Cardiomyocyte Microtubules: Control of Mechanics, Transport, and Remodeling. Annu Rev Physiol 84:1–27. doi: 10.1146/annurev-physiol-062421-040656

36. Webster DR, Borisy GG (1989) Microtubules are acetylated in domains that turn over slowly. J Cell Sci 92 (Pt 1):57–65

37. Webster DR, Wehland J, Weber K, Borisy GG (1990) Detyrosination of alpha tubulin does not stabilize microtubules in vivo. J Cell Biology 111:113–122. doi: 10.1083/jcb.111.1.113

38. Xu Z, Schaedel L, Portran D, Aguilar A, Gaillard J, Marinkovich MP, Théry M, Nachury MV (2017) Microtubules acquire resistance from mechanical breakage through intralumenal acetylation. Science 356:328–332. doi: 10.1126/science.aai8764

