## Supplementary material for "Desmin intermediate filaments and tubulin detyrosination stabilize growing microtubules in the cardiomyocyte": All supplemental figures

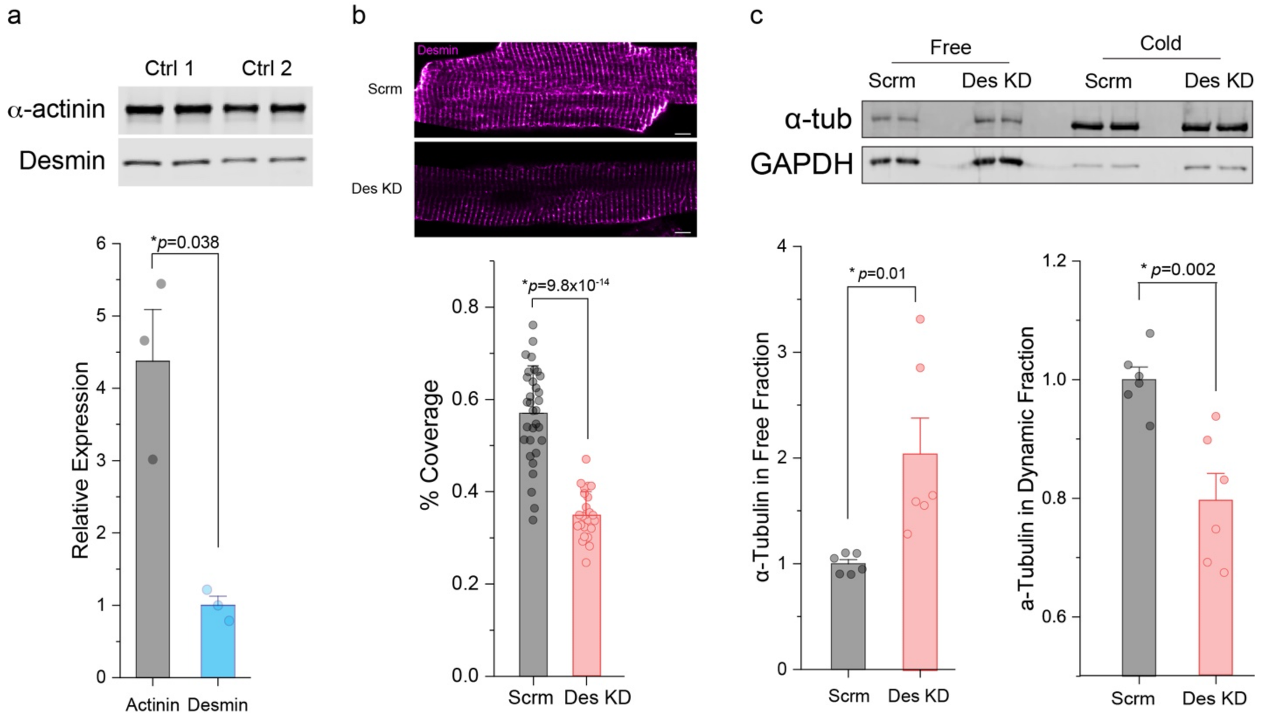


**Supplemental Fig. 1 (a)** Representative western blot images **(top)** and quantification **(bottom)** of α-actinin and desmin using whole cell lysates from adult rat cardiomyocytes; the blots were immunostained using the same α-actinin and desmin antibodies and concentration as used in Figure 1e (N=3 rats, n=3 averaged-values from 2 WB technical lanes per rat). **(b)** Representative immunofluorescent images **(top)** and quantification of coverage area **(bottom)** for desmin in scramble shRNA (Scrm) and Desmin Knock-Down shRNA (Des KD) adult rat cardiomyocytes (N=3 rats, n=~10 cells per rat). **(c)** Representative western blot **(top)** and quantification **(bottom)** of α-tubulin and GAPDH in free and cold-sensitive fractions from rat cardiomyocytes infected with adenovirus containing Scrm and Des KD (N=3 rats, n=6 WB technical lanes). Bar represents mean ±1 SEM; statistical significance determined with Two-sample Student's T-test.

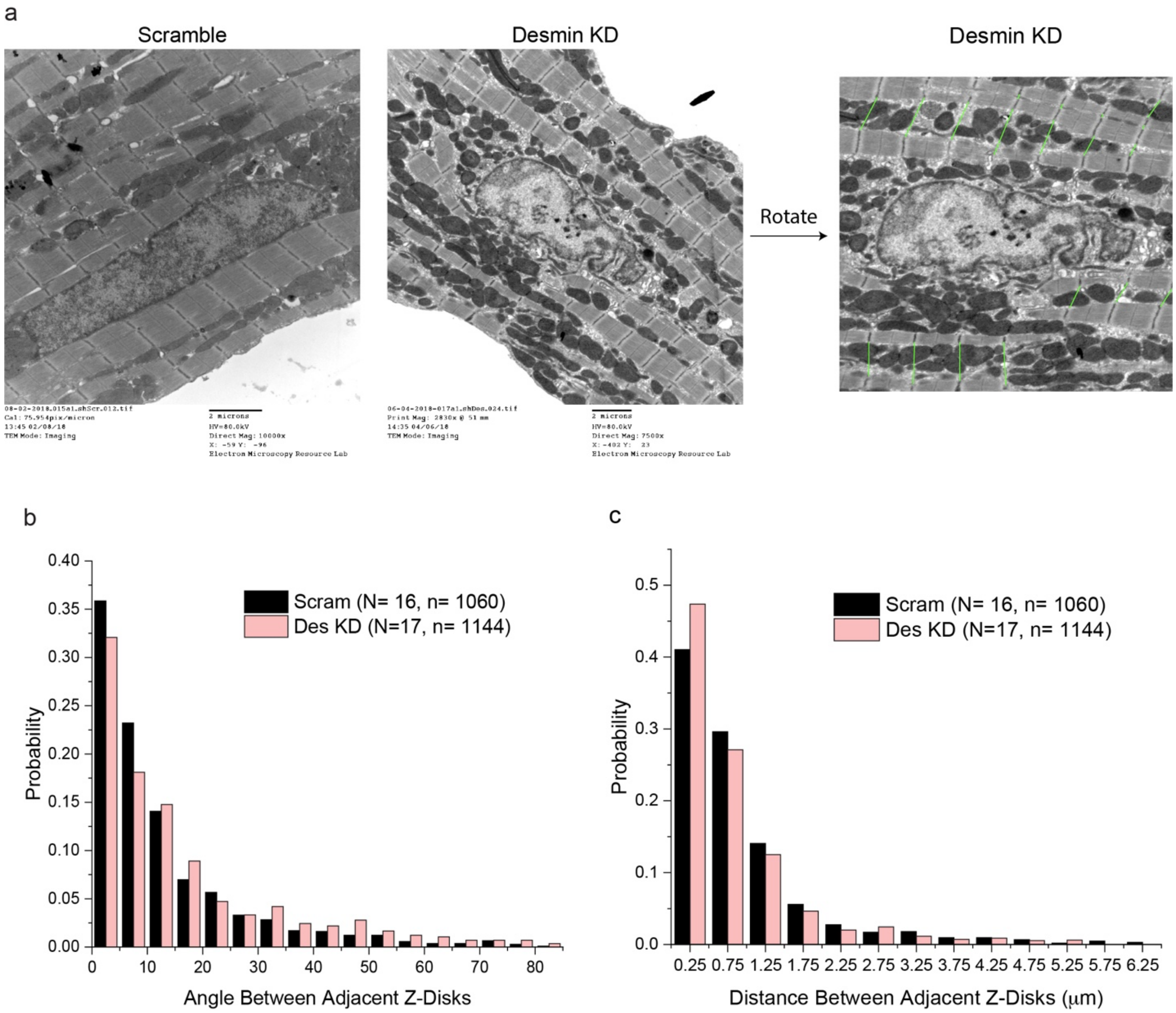


**Supplemental Fig. 2 (a)** Representative electron microscopy images of isolated rat cardiomyocytes with or without desmin knockdown. **(b)** Histograms of angle and **(c)** distance between adjacent Z-discs with or without desmin knockdown (N=cells, n=events).

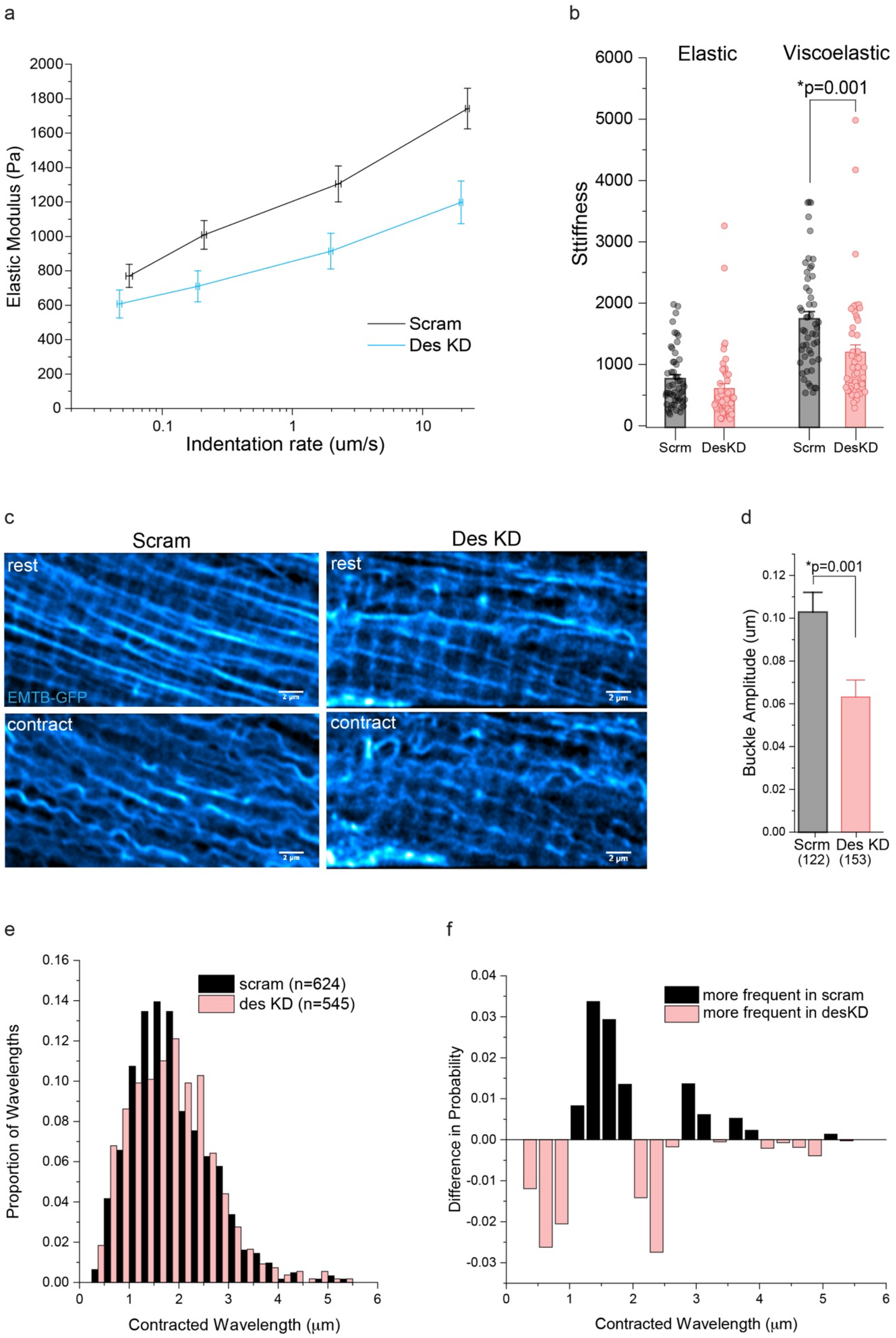

a
Elastic Modulus (Pa)
Indentation rate (um/s)
Scram
Des KD
b
Elastic
Viscoelastic
*p=0.001
Stiffness
Scrm
DesKD
Scrm
DesKD
c
Scram
Des KD
rest
rest
EMTB-GFP
contract
contract
2 μm
d
*p=0.001
Buckle Amplitude (um)
Scrm
(122)
Des KD
(153)
e
scram (n=624)
des KD (n=545)
Proportion of Wavelengths
Contracted Wavelength (μm)
f
more frequent in scram
more frequent in desKD
Difference in Probability
Contracted Wavelength (μm)

**Supplemental Fig. 3 (a)** Nanoindentation measurement of cardiomyocyte viscoelasticity displayed as stiffness (elastic modulus) as a function of probe indentation velocity with or without desmin KD. **(b)** Quantification of rate-dependent (viscoelastic) and independent (elastic) stiffness; height of bar graphs denotes mean and error bar ±1 SE. **(c)** Representative images of microtubules in a cardiomyocyte at rest **(top)** and at the peak of contraction **(bottom)** with or without desmin KD. **(d)** Quantification of microtubule buckle amplitude; bar represents mean ± 1 SEM (n=cells). **(e)** Histogram of wavelength distribution of individual microtubule buckles. **(f)** distribution difference (scram – des KD from E). Statistical significance determined using Two-sample Student's T-test.

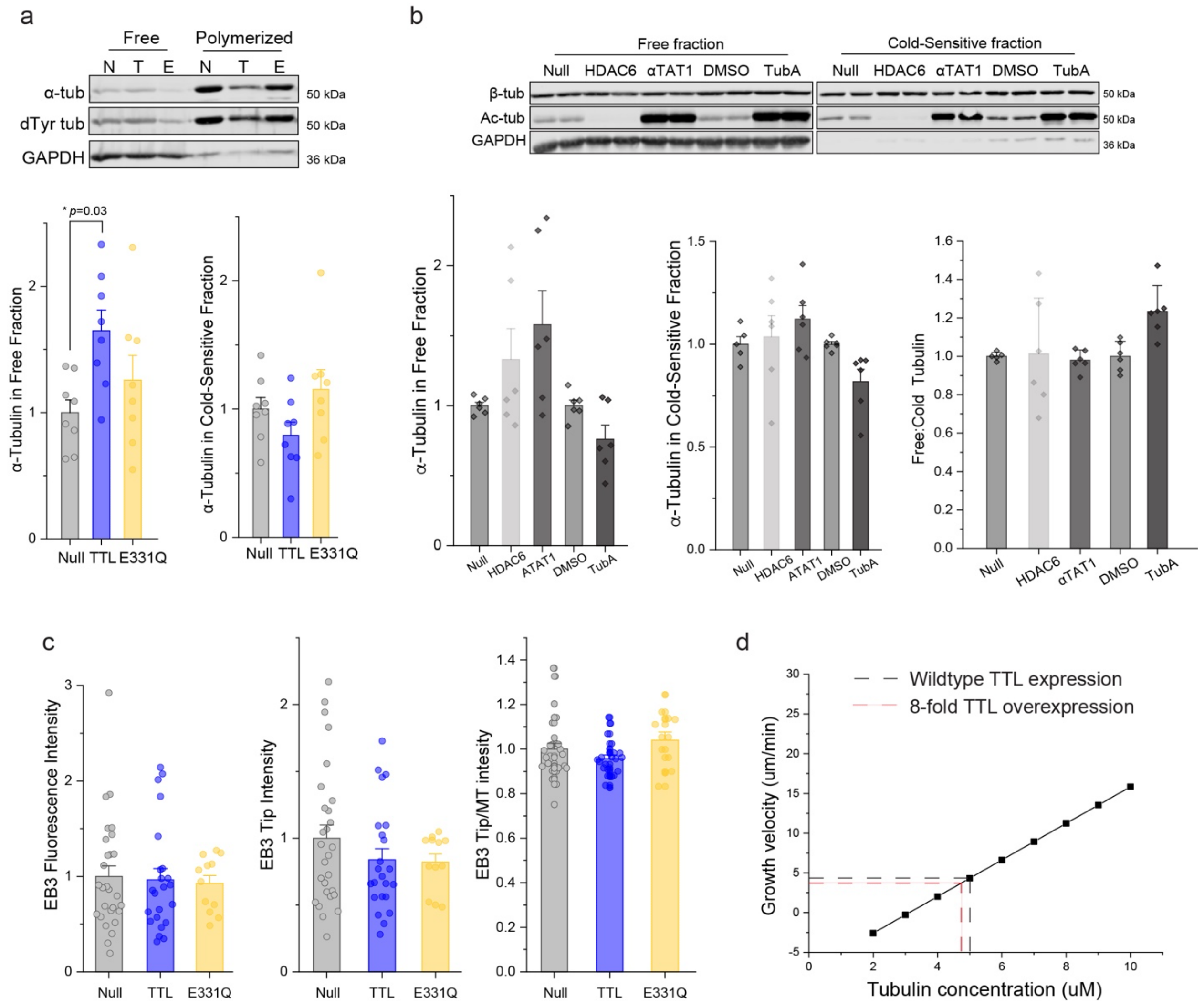


**Supplemental Fig. 4 (a)** Representative western blot **(top)** and quantification **(bottom)** of a-tubulin in free and cold-sensitive fractions from adult rat cardiomyocytes infected with null, TTL, or E331Q adenovirus (N=4 rats, n=8 WB technical lanes). **(b)** Representative western blot **(top)** and quantification **(bottom)** of β-tubulin and acetyl tubulin in free and cold-sensitive fractions from adult rat cardiomyocytes infected with either null, HDAC, or ATAT1 adenovirus, or treated with DMSO or TubA (N=3 rats, n=6 WB technical lanes). **(c)** Background **(left)** EB3-GFP fluorescence intensity, EB3 tip fluorescence intensity **(middle)**, and the ratio of EB3 tip fluorescence intensity to microtubule EB3 intensity **(right)** in adult rat cardiomyocytes co-infected with EB3-GFP and either Null, TTL, or E331Q adenoviruses. Bar represents mean ± 1 SEM; statistical significance determined using one-way ANOVA with post-hoc test. **(d)** To reduce levels of detyrosination in this study we relied on overexpression of TTL, which is known to associate with the free α/β tubulin heterodimer in a 1:1 complex with a $K_d$ of 1 μM[36]. Due to the nature of this interaction, TTL overexpression can decrease the amount of "polymerization-competent" tubulin, leading to a decrease of *in vitro* microtubule polymerization[36]. The relationship between free tubulin concentration and microtubule polymerization is described as a simple 1D model below, where $\nu_g$ is the velocity of microtubule growth, *d* is the dimer length (assumed to be 8 nm), $k_{on}$ is the tubulin dimer association constant per protofilament (assumed to be 4.8 $\mu M^{-1}s^{-1}$ )[24], $k_{off}$ is the tubulin dissociation rate constant per protofilament (assumed to be $15s^{-1}$ )[24], and [Tb] is the concentration of free tubulin[4].

$$\nu_g = d\left(k_{on}[Tb] - k_{off}\right).$$

Solving for the equation of the line at a physiologic tubulin concentration of 5μM provides a theoretical value of 4.32 $\mu m\ s^{-1}$ for growth velocity. Using the $K_d$ value for TTL-tubulin interaction at a tubulin concentration of 5μM suggests that an 8-fold overexpression of TTL[9] will produce 0.2 μM of TTL-tubulin complex. If we assume that tubulin bound to TTL cannot be polymerized, then only 4.8 μM of free tubulin is polymerization-competent at any time. This change in polymerization-competent tubulin would lead to ~10% decrease in growth velocity. We did not observe an effect of E331Q on microtubule growth kinetics or on event frequency suggesting that the local tubulin concentration at the microtubule plus-tip is unaffected by the sequestration activity of TTL. Bar represents mean ± 1SEM; statistical significance for (a) and (c) was determined using one-way ANOVA with post-hoc test and for (b) was using Two-sample Student's T-test.

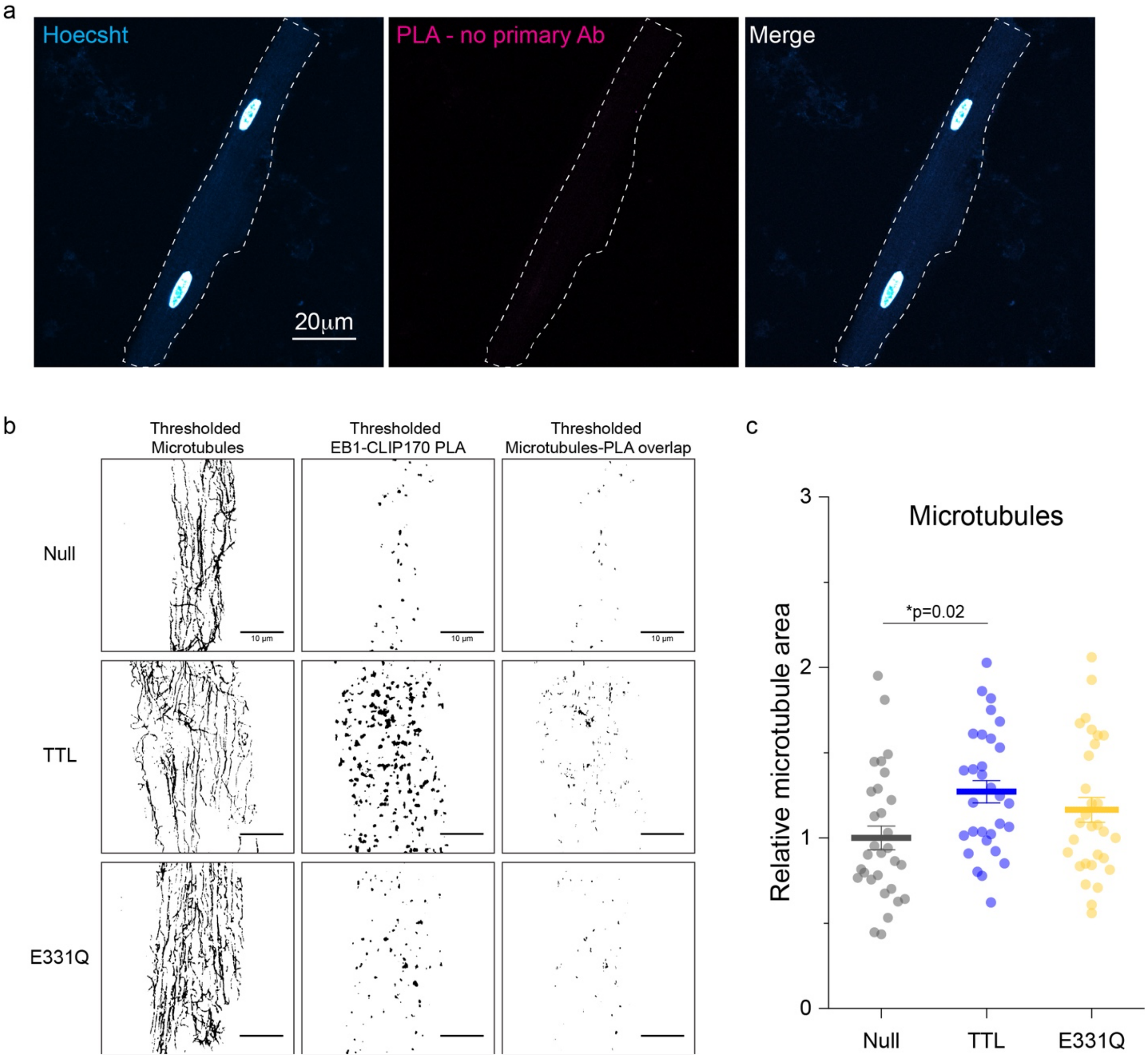


**Supplemental Figure 5. (a)** Representative negative control immunofluorescent images after PLA in adult rat cardiomyocytes without EB1 or CLIP170 primary antibodies. **(b)** Representative thresholded immunofluorescent images used in the analysis and quantification of EB1-CLIP170 in Figure 5d. **(c)** Quantification of microtubule area in adult rat cardiomyocytes treated with Null, TTL, or E331Q adenovirus for 48h (N=3 rats, n=10 cells per rat). Bar represents mean ±1 SEM; statistical significance determined with one-way ANOVA with post-hoc test.
